# Development of new food-sharing relationships among nonkin vampire bats

**DOI:** 10.1101/534321

**Authors:** Gerald G. Carter, Damien R. Farine, Rachel J. Crisp, Julia K. Vrtilek, Simon P. Ripperger, Rachel A. Page

## Abstract

Social relationships that involve costly helping occur most often among kin, but in many complex and individualized animal societies, nonkin also demonstrate stable cooperative relationships that share similarities with human friendship. How do such cooperative bonds form between complete strangers? Here, we show evidence that unfamiliar nonkin vampire bats (*Desmodus rotundus*) selectively escalate low-cost investments in allogrooming before developing higher-cost food-sharing relationships. By introducing female bats from geographically distant sites in captive pairs or groups and fasting them repeatedly over 15 months, we observed that increasing rates of grooming a partner predicted the occurrence of that partner’s first food donation back to the groomer, after which grooming rates no longer increased. New food-sharing relationships formed in 14.5% of 608 possible female pairs, emerged in a reciprocal pattern, and developed more often when strangers lacked alternative familiar partners. Our results are consistent with predictions from the ‘raising-the-stakes’ hypothesis that strangers ‘test the waters’ of new relationships by initially making low-cost investments in grooming before making higher-cost food donations. This form of ‘raising the stakes’ (e.g., transitions from clustering to grooming to food sharing) might play an underappreciated role in many other social decisions with long-term consequences, such as joining a new social group or choosing a long-term mate.

**Significance statement:** Vampire bats form long-term cooperative social bonds that involve reciprocal regurgitated blood sharing. But how do two individuals go from complete strangers to reciprocal food donors? By introducing unfamiliar bats, we found evidence that low-cost grooming paves the way for higher-cost food donations. Bats that formed new food-sharing relationships had a history of escalating reciprocal grooming up until the food sharing began. Food sharing emerged in a reciprocal fashion and it emerged more often when two strangers could not access their original groupmates. The finding that unfamiliar nonkin vampire bats appeared to gradually and selectively transition from low-cost to high-cost cooperative behaviors is the first evidence that nonhuman animals ‘raise the stakes’ when forming new cooperative relationships.

## Text

Animal societies are fundamentally shaped by repeated interactions among individuals over time. Repeated interactions allow individuals to choose to cooperate based on their past experience across different partners (1–3). Organisms as diverse as animals, plants, and fungi have demonstrated partner choice: individuals prevent exploitation by shifting their cooperative investments towards partners that provide better reciprocal returns (3–7). Across several nonhuman mammals, repeated cooperative interactions lead to adaptive and enduring social bonds that share similarities with human friendship (8–12), but it remains unclear how these initially form. A significant challenge has been understanding how individuals prevent exploitation while forming these stable bonds. How do complete strangers develop a long-term cooperative relationship?

A key idea is that individuals should reduce the risk of exploitation by initially spreading out smaller cooperative investments across time (‘parceling’ (13)) or across different partners (‘social bet-hedging’ (14)), and then gradually escalating investments in the most cooperative partnerships (‘raising the stakes’ (15)). For example, one might first assess a potential partner’s tolerance by clustering for warmth, then gain feedback by grooming the partner, and then use the partner’s response to decide whether to provide higher-cost food donations or coalitionary support (16). Despite its intuitive appeal for explaining how new cooperative relationships develop, evidence supporting the 20-year-old ‘raising-the-stakes’ model (15) is surprisingly scarce. An early test using the cleaner and client fish mutualism suggested that the model does not apply well to situations with severe asymmetries in partner payoffs or options (17). Studies with nonhuman primates (18–21) have tested only snapshots of established relationships rather than the formation of new ones. Human strangers ‘raise the stakes’ when making monetary bids in cooperation games (e.g. 22, 23), but we currently lack supporting evidence for this strategy in the more ecologically-relevant context of relationship formation. Gathering this evidence requires measuring the emergence of natural helping behaviors between randomly introduced strangers.

We tracked the development of new cooperative relationships between previously unfamiliar wild-caught vampire bats (*Desmodus rotundus*) over 15 months. Cooperative relationships in this species involve low-cost allogrooming (hereafter *grooming*) and higher-cost regurgitations of ingested blood, or *food sharing* (14, 24–28). We found support for several lines of evidence that vampire bats use reciprocal grooming to gradually establish new bonds that include food donations. If bats choose partners, then new food-sharing relationships should form more often when bats have fewer alternative partners. If grooming gradually leads to sharing, then grooming rates should predict the probability that the grooming recipient later donates food back to the groomer. Grooming rates should also start low and increase over time but only up until the first reciprocal food donation. Finally, new food sharing should be rarer, occur after mutual grooming, and emerge in a reciprocal fashion.

Female vampire bats demonstrate kin-biased fission-fusion social dynamics (24, 27, 28). New nonkin social bonds can form when an unrelated female joins a social network about once every two years (24, 28), with individual bats living for up to 16 years in the wild (29). To observe how new food-sharing relationships form between adults, we captured adult females from two distant sites in Panamá, Tolé (n=19) and Las Pavas (n=8), and we then ran 638 fasting trials in which an overnight-fasted subject could be fed by a previously unfamiliar bat from another site. To test the prediction that new sharing relationships would form more often when strangers have fewer partner options, we compared the occurrence of new sharing when wild-caught strangers were introduced in isolated pairs (one Las Pavas and one Tolé bat), in small groups (one Las Pavas and three Tolé bats), or in one large mixed group (all bats together; see Methods, Supplementary Information (SI) Appendix, Fig. S1). New bonds can also form when individuals are born into a group, and these relationships might form differently. We therefore also measured the development of non-maternal cooperative relationships between 26 female adults and 13 younger captive-born bats (6 males and 7 females, 11 to 21 months old) in the large mixed group.

To test our hypotheses, we compared the observed coefficients from general and generalized linear models (slopes β, and odds ratios OR, respectively) to expected distributions of coefficient values expected under the null hypotheses using permutations of the network or the event data (see Methods). We use the term ‘potential relationship’ for a pair of bats that could have groomed or shared food, the word ‘relationship’ for an *observed* network edge (directed), and the word ‘bond’ to discuss the underlying construct that we *inferred* from the observed relationship (see SI, Table S1).

Fasted bats were fed by at least one donor in 61% of trials (SI Appendix 2). Over 424 days and 12,012 opportunities for new food donations, new food sharing developed in 10.8% of the 996 potential relationships among all bats, 14.5% of 608 potential relationships among females, and 15.6% of 243 potential relationships among wild-caught adult females (SI Appendix 3). All bats had at least one donor (range=1-16, mean=6.6). The average number of new food donors per adult female bat was 2.7 (range=0-7) and the average per captive-born bat was 2.6 (range=0-6). New grooming relationships developed far more frequently (all bats=51.9% of 1008; females=58.9% of 618; wild-caught adult females=78.2% of 248). The average number of new groomers was 7.2 (range=0-16) for adult females and 14.4 (range=1-23) for captive-born bats.

If bats choose new partners based on their phenotype alone, then relationships should form more often when bats have more alternative partners. On the other hand, if bats ‘test the waters’ of each new relationships, they should choose partners based on both the availability of different partners and their past experiences with each, and food-sharing relationships should therefore form more often when bats have *fewer* alternative partners. As expected, when strangers from Las Pavas and Tolé were introduced and housed as isolated pairs, we observed higher rates of new food sharing (β=1.14, p=0.002) and new grooming (β=1.09, p=0.02) compared to when one Las Pavas bat was introduced to three Tolé bats, despite there being fewer potential new bonds available to form (SI Appendix 4). When we aggregated bats from the controlled introduction trials into a large mixed group, bats preferentially fed and groomed their original familiar groupmates, and new sharing emerged even more gradually than in the isolated pairs or in small groups (SI Appendix 5, Fig. S2 and S3).

If the bats use low-cost grooming to build higher-cost sharing bonds, then the grooming rate should predict the probability of the first food donation in the opposite direction. As expected, new food sharing emerged on days after we observed mutual grooming more than expected by chance (SI Appendix 6), and the grooming rate given by actor A to recipient B predicted the later occurrence of new food sharing from B back to A (OR=2.15, p=0.0002, n=897). The trajectory of grooming rates over time clearly differed between pairs that developed new food-sharing relationships versus pairs that did not (interaction: OR=1.60, p<0.0001, Fig. 1). The slope of this increase in grooming was also greater *before* the first reciprocal food donation than after. Initial grooming rates started low, then increased over time up until the new food-sharing relationship formed (Fig. 2).

**Fig. 1.**
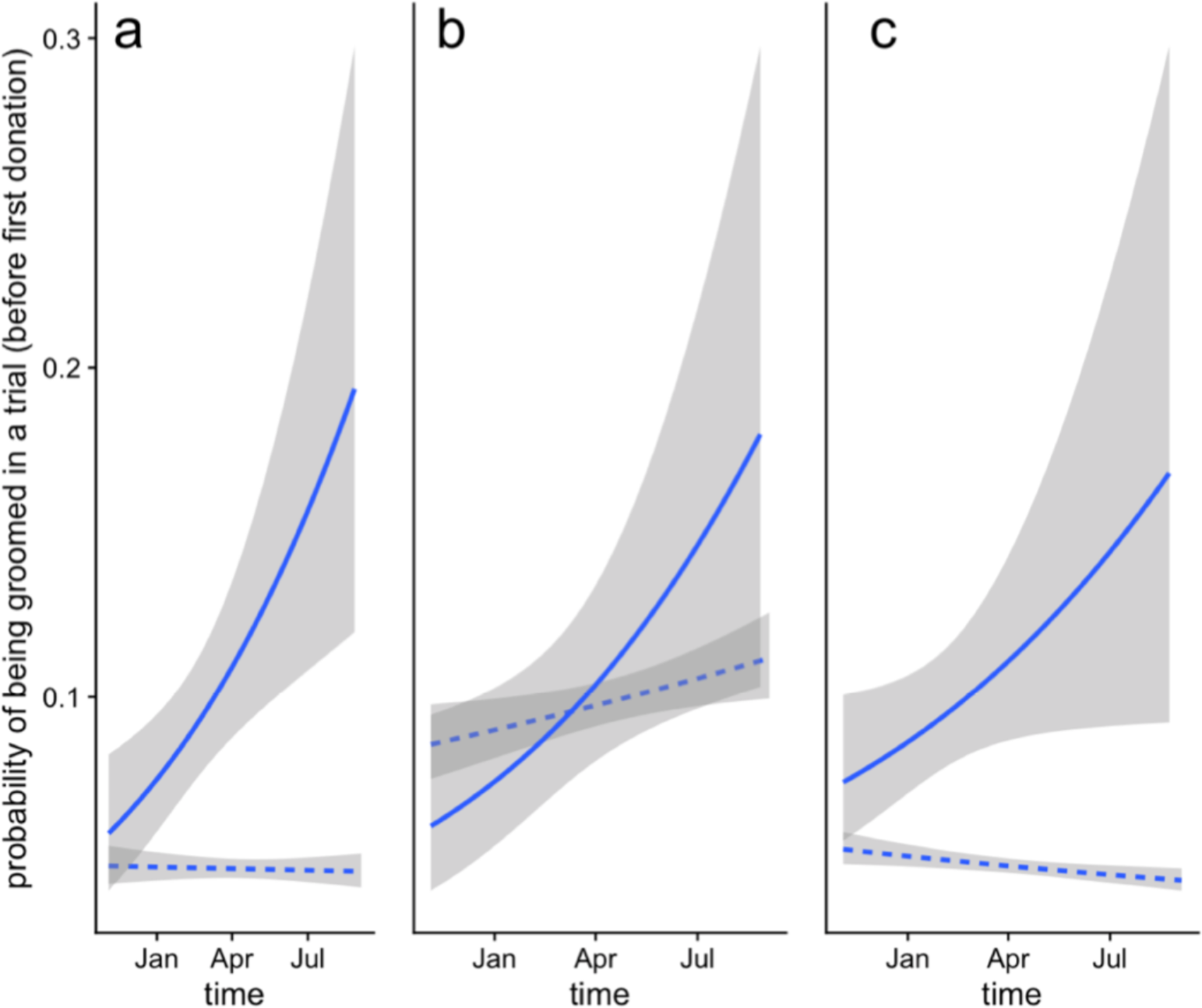
Increasing A-to-B grooming led to new B-to-A food-sharing relationships. In cases where a new food-sharing relationship formed (solid line), the grooming rate towards the future donor increased over time before the first donation occurred (OR=1.40, n=33, p<0.0001), but the grooming rate towards a potential donor remained low in cases where no food-sharing relationship formed (dashed line; OR=0.99, n=420, p=0.58). This divergence in all potential new relationships (panel a) was also detected within previously unfamiliar adults (panel b), and within relationships with captive-born bats (panel c), which had more divergent grooming trajectories (SI Appendix 7). Shading shows the 95% CI for the fitted model’s predictions.

**Fig. 2.**
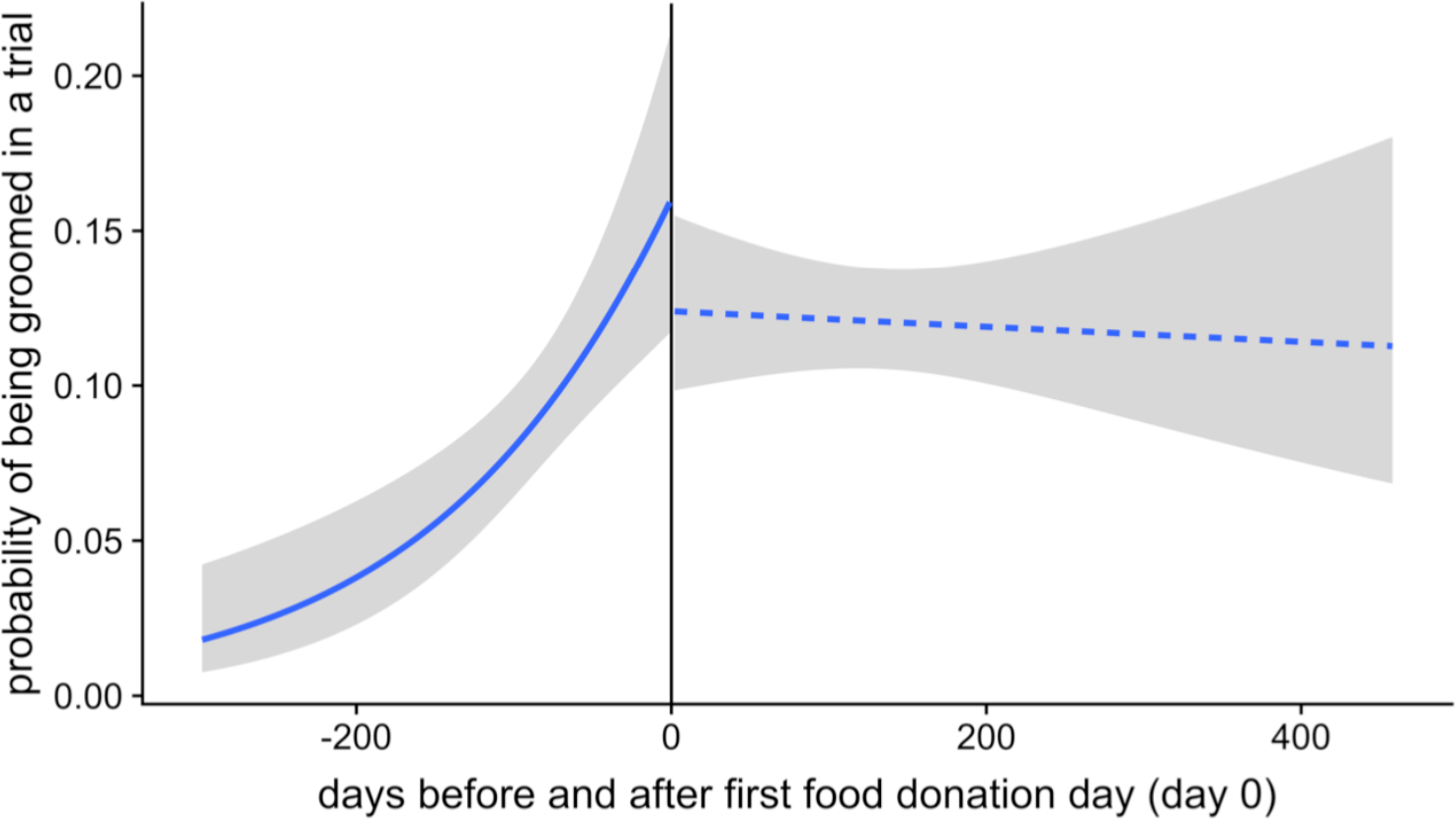
Grooming rates increased before, but not after, new food-sharing occurred. The probability of a focal bat grooming the new donor in a 1-h trial (y-axis) increased before the first day that the donor fed the focal bat (i.e. ‘day zero’; OR=1.4, p=0.0005), but not after this day zero (OR=1.01, p=0.47; interaction: OR=1.57, p=0.0003). This effect was seen in new food-sharing relationships with or without captive-born bats (SI Appendix 8). Shading shows the 95% CI for the fitted model’s predictions.

Emergence of new food sharing was more reciprocal than expected by chance, even when controlling for kinship (MRQAP-DSP; reciprocal sharing: β=0.33, p<0.0002, kinship: β=0.02, p=0.65; SI Appendix 9). Grooming rates in new relationships were also symmetrical across dyads but we lacked the power to determine whether grooming symmetry increased over time within dyads (see SI Appendix 10). Grooming rates were highest between bats that formed two-way food-sharing relationships, intermediate in relationships where we observed sharing in only one direction, and lowest in pairs where we never saw food sharing (SI, Fig. S4).

The rarity of new food-sharing relationships corroborates past evidence that food regurgitations are energetically costly and that food-sharing bonds require investments of time and energy (14, 24, 30, 31). The relationship between new grooming and new food sharing was unlikely to be caused by mere proximity because the effect of new grooming on new food sharing remained evident regardless of whether or not bats were forced into close proximity (SI Appendix 11).

These findings provide the clearest evidence to date that nonkin food sharing in vampire bats is not a byproduct of kin selection (26). Before this study, one hypothesis was that food sharing decisions among nonkin could depend entirely on heuristics based on phenotypic similarity, resulting in a spurious pattern of symmetrical helping that looks like reciprocity (32, 33). However, this hypothesis incorrectly predicts that food-sharing relationships should form immediately and occur most frequently in larger groups simply because there are more opportunities for similar matching phenotypes.

Our results were consistent with the hypothesis that relationships formation occurs through some form of ‘raising the stakes’ (15). This model has yet to be tested during the transition from ‘strangers’ to ‘friends’ because these changes are difficult to document in nature (see SI Appendix 12). Past evidence for the ‘raising-the-stakes’ strategy (15) has also been scarce in part because it is a variation on the classic ‘tit-for-tat’ strategy in the iterated prisoner’s dilemma (1), a model which is difficult to test using natural forms of cooperation (4, 32, 34). ‘Tit-for-tat’ forms of reciprocity are demonstrated by experiments with trained instrumental tasks and payoffs that accrue in distinct rounds, such as rats taking turns to pull a lever to deliver food (4–6). However, the ‘tit-for-tat’ model excludes many factors crucial in the real world, including partner choice, partner fidelity, exchange of multiple service types, and the many cost-benefit asymmetries resulting from demography, market effects, and social rank (4, 7, 34). If social bonding involves integrating many different kinds of social interactions into a single positive association, one should not expect clearly alternating exchanges of help. In primates, cooperation within long-term social bonds does not produce strict ‘tit-for-tat’ exchanges of help; strongly bonded partners show *less* evidence for short-term contingencies in grooming (9).

To clearly demonstrate that an actor’s cooperative investments are contingent on a partner’s previous behavior, one must prevent reciprocation and then detect a subsequent decrease in the actor’s cooperative investment. This evidence of reciprocity has yet to be experimentally demonstrated in food-sharing vampire bats or in any other long-term social relationship (SI Appendix 13). Our findings show that such an experiment would be most powerful if researchers targeted newly developing relationships rather than established ones, and if they tracked multiple cooperative behaviors rather than just one. Past studies on ‘raising the stakes’ during relationship development have focused on increasing rates of a single cooperative behavior (18–23), but individuals can also raise the stakes by adding new higher-cost behaviors. Our findings suggest that female vampire bats do both, first increasing grooming rates and then transitioning from low-cost grooming to high-cost food-sharing.

The relevance of our findings extends beyond high-cost cooperative behaviors. For example, in some species, courtship behaviors could be seen as a short-term investment in the formation of longer-term pair bonds with substantial fitness consequences (35). Similarly, the role of mere physical contact as a low-cost method for building tolerance and trust might be more general than currently recognized. The key role of grooming for relationship maintenance in primates is well established, but growing evidence suggests that similar tactile behaviors can reduce fear and encourage tolerance and cooperation in many other species of mammals, birds, and fish (e.g. 5, 36–42). Recently developed methods for tracking formation of social bonds at fine temporal scales (43, 44) could provide new opportunities to test whether gradual escalation of proximity and body contact is a widespread mechanism for socially ‘testing the water’.

## Methods

### Animals

We conducted experiments at the Smithsonian Tropical Research Institute in Gamboa, Panama. We used 41 common vampire bats (*Desmodus rotundus*) as subjects, including 19 female bats captured exiting a roost in Tolé, Panamá; 8 female bats captured foraging at a cattle pasture in Las Pavas, Panamá about ~215 km from Tolé; and 14 captive-born bats (8 females, 6 males). We studied adult females and their young, because these individuals form the basis of food-sharing networks in the wild, whereas adult males compete for access to territories and females and do not form stable bonds as often (24–28). To ensure familiarity within groups and unfamiliarity between groups, we housed the groups separately (Tolé bats for 6 months and Las Pavas bats for 2 weeks) before the study began. Bats were marked with subcutaneous passive integrated transponders (Trovan Ltd. USA) and a visually unique combination of forearm bands (Porzana, National Tag, and birdbands.com). To feed bats, we provided refrigerated or thawed cattle or pig blood defibrinated with sodium citrate and citric acid.

We used a 3-4 mm biopsy punch to collect tissue samples in 80% or 95% ethanol, then used a salt–chloroform procedure for DNA isolation, and a LI–COR Biosciences® DNA Analyser 4300 and the SAGA GT allele scoring software to genotype individuals at 17 polymorphic microsatellite loci. Allele frequencies were based on 100 bats from Tolé and 9 bats from Las Pavas, respectively. Genotypes were 99.9% complete. To estimate genetic relatedness, we used the Wang estimator in the R package ‘related’. To estimate kinship, we assigned a zero kinship to known unrelated individuals from different sites and to individuals with negative pairwise relatedness, and we assigned a kinship of 0.5 for known mother-offspring pairs or pairs with genetic relatedness estimates greater than 0.5. For all other pairs, we used genetic relatedness as the estimate for kinship.

### Experimental design

We induced allogrooming and regurgitated food sharing using a fasting trial, in which a focal subject was isolated from the group without food for a night and a day, then released back to the group of fed bats for 1 hour the following night. During the hour, all grooming or food-sharing interactions with the subject were recorded using an infrared (IR) light and an IR-sensitive video camera. Each food sharing bout was estimated by the number of seconds that the unfed subject spent licking the mouth of a particular groupmate. Grooming was defined as chewing or licking the fur or wings of another bat. The dyadic sharing or grooming for a trial was estimated as the sum of all bouts that were at least 5 seconds long. We weighed bats before and after trials. Observed mouth-licking durations predicted weight gain (SI Appendix 1).

We conducted fasting trials in each group during three experimental phases (SI, Fig. S1). First, we conducted 57 ‘baseline’ trials to assess preliminary sharing rates between the 19 Tolé bats housed in a 1.7 × 2.1 × 2.3 m outdoor flight cage (3,420 possible sharing interactions in one group). Second, we conducted 106 ‘controlled introduction’ trials to assess possible formation of new food-sharing bonds between bats introduced as either an isolated pair (one Las Pavas bat and one Tolé bat) or a quartet (one Las Pavas bat and three Tolé bats), housed in a 28 × 28 × 40 cm clear plastic observation cage (10 pairs and 8 quartets). These controlled introductions provided for 162 opportunities for new food sharing between previous strangers (SI, Table S2). Finally, we conducted 532 ‘mixed-group’ trials to assess the formation of new sharing relationships when all bats were housed together in the flight cage described above (19 Tolé, 7 Las Pavas, and 14 captive-born bats). The introductions in this combined group provided 11,823 more opportunities for new sharing.

### Statistical analyses

During the baseline and mixed-group trials, we estimated food donation size as the number of seconds that a fasted subject spent mouth-licking a fed groupmate. During the controlled introduction trials, however, when bats were forced in close proximity, we saw a greater frequency of begging, defined as mouth-licking that is clearly not food-sharing because the partner is turning away from the mouth-licking bat and the mouth-licking bat does not gain the weight that would be expected from food-sharing. To be conservative when measuring sharing, we therefore did not count mouth-licking as food sharing during the controlled introduction trials unless the subject weighed more than expected based on the average weight change for bats that did not perform any mouth-licking.

Durations of sharing and grooming were lognormal. To create a standard index of grooming rates, we therefore transformed the total duration of directed dyadic interactions in each trial using natural log (x+1). We call these measures of the log duration per hour ‘rates’. When interaction bout duration and probability had different meanings, we decomposed rates into two separate response variables: amounts (the magnitude of nonzero rates in a trial) and probabilities (the presence or absence of a nonzero rate in a trial). We used permutation tests with 10,000 permutations for p-values and bootstrapping for all 95% confidence intervals. Null distributions were not always centered on zero due to structure in the data, so caution must be taken when considering the observed coefficients.

Grooming could occur before sharing simply because it is more frequent. To test whether mutual grooming preceded new sharing more than expected by chance, we compared the observed probability of observing mutual grooming before new sharing to the values expected from a null model based on randomly swapping the label of interactions (grooming versus sharing) within each dyad. This permutation test controls for the relative frequency and timing of grooming and sharing events in new dyads. To test for ingroup-outgroup biases in sharing for each site, we calculated observed coefficients for the effect of the actor and receiver being from the same capture site on actor grooming rates, then we calculated expected coefficients by permuting the grooming rates within each actor to different possible recipients.

To test the effects of kinship and reciprocal grooming on the formation of new food-sharing relationships in the mixed-group trials, we used multiple regression quadratic assignment procedure with double semi-partialing (MRQAP-DSP) via the netlogit function in the sna R package. We also used this method to test the effect of grooming on occurrence of new sharing only within the controlled introduction trials. This procedure uses generalized linear models via the glm function in lme4 package to calculate the observed coefficients and uses network-level permutations to get expected coefficients. Since MRQAP-DSP cannot test interaction effects, we compared observed and expected interaction coefficients using permutations in which we shuffled trial rates given by the actor among different possible receivers and then shuffled the trial rates received by the receiver among different possible actors. If the interaction coefficients were significant (p<0.05), we conducted separate MRQAP-DSP tests within each group.

To test whether interaction rates changed over time, we generated expected coefficients for general or generalized linear models by permuting the order of interactions within each potential relationship. One captive-born bat died for unknown reasons during the mixed-group trials, so we removed it from all temporal analyses. To test for evidence of reciprocal sharing, we used MRQAP-DSP to test if the matrix of new sharing in the mixed-group trials was predicted by reciprocal sharing when controlling for kinship. As an additional test, we also counted the occurrence of both novel sharing and reciprocal sharing for all new potential relationships, then counted the same number after randomizing the presence of sharing across potential relationships.

## Data availability

Behavioral data, genotypes, and R code are available as supplementary information.

## Acknowledgments

We thank the Smithsonian Tropical Research Institute for logistical support. Isabelle Waurick conducted the molecular lab work. Frieder Mayer enabled the molecular lab work. Work was supported by a Smithsonian Postdoctoral Fellowship, a Humboldt Research Fellowship, a Smithsonian Institution Scholarly Studies Grant and a grant from the National Geographic Society Committee for Research and Exploration (WW-057R-17).

## Author contributions

Conceptualization, GC; Methodology, GC; Investigation, GC, RC, JV; Genotyping, SR; Statistical analysis, GC; Original draft, GC; Review & Editing, GC, RC, JV, SR, DF, RP; Funding acquisition, GC, SR, RP; Resources, RP; Supervision, GC, DF, RP;

## Notes

#### Summary of Updates

Revised for clarity and added new supporting analyses.

