## Supplementary material for "Development of new food-sharing relationships among nonkin vampire bats": SI

**Supplementary Information**

For the paper, Carter et al. *Development of new food-sharing relationships among nonkin vampire bats.*

**Appendix**

*1. Evidence of food sharing*

Fasted subjects gained an average of 51 mg of mass per minute of mouth-licking ($R^2$ = 0.75, p < 0.001, 95% CI: 45 to 57 mg/min, n = 619 trials without missing data), which is comparable to previous estimates from another captive colony (38 mg of mass per minute of mouth-licking, $R^2$ = 0.67, 95% CI: 33 to 46 mg/min, n = 121 trials, colony described in [1-3]).

*2. Rates of food sharing*

Across all trials, the probability that a given bat received food from any groupmate was 61% (95% CI = 57 to 64%, 41 bats, 693 trials), which is much lower than the 95% success rate observed in the previous long-term captive colony (95% CI = 92 to 98%, 29 bats, 183 trials; described in [1-3]). Assuming that mouth-licking events over 5 seconds were food donations, 64% of the 340 mixed-group trials with food sharing involved one donor, 24% had two donors, 9% had three donors, 2% had four donors, and two trials had up to five donors (i.e. mouthlicking durations of 5 s or longer).

*3. Development of new food-sharing relationships*

We induced 12 of the 38 new food-sharing relationships between wild-caught adult females during the 106 'controlled introduction' fasting trials (Table S2), in which a single stranger Las Pavas bat was introduced to either one unfamiliar Tolé bat (forming an isolated pair) or to three Tolé bats (forming a quartet). The 26 other new food-sharing relationships between wild-caught adult females developed gradually during the 532 'mixed-group' fasting trials in the subsequent period when all the bats from both groups could freely interact (see Figures S1, S2, S3).

During the mixed-group period, there was a 10% chance that a new food-sharing relationship would develop between an adult and a captive-born bat (7 females, 6 males, 3-19 months old; 68 of 748 potential relationships) and 3.9% chance of a new food-sharing relationship between two captive-born bats. Captive-born bats groomed and fed each other less than they groomed and fed adult females that were not their mother (grooming: $\beta$ = -0.09, n = 463, p = 0.009; sharing: $\beta$ = -0.16, n = 463, p < 0.0001).

During the mixed-group trials, the Las Pavas bats were biased towards feeding and grooming other Las Pavas bats (sharing: $\beta$ = 0.28, n = 160, p < 0.0001, grooming: $\beta$ = 0.53, n = 160, p < 0.0001). Tolé bats showed the same within-group bias for sharing ($\beta$ = 0.09, n = 390, p = 0.003) but not grooming ($\beta$ = 0.10, n = 390, p = 0.12). New sharing relationships with captive-born bats were also more likely among bats from the same source population (OR = 1.73, p = 0.04). After controlling for this within-group bias, we found no evidence for a kinship bias in grooming (MRQAP-DSP, $\beta$ = 0.12, p = 0.57) or sharing (MRQAP-DSP, $\beta$ = 0.21, p = 0.15). We lacked the statistical power to test for increases in food donation sizes over time within new dyads, but when pooling donations across all dyads, new donations that occurred later in time were not significantly larger ($\beta$ = 5.6, n = 37, p = 0.94).

*4. Food sharing relationships emerged faster in isolated pairs than in quartets*

The seven food donations in new relationships in isolated pairs tended to occur sooner on average (mean latency = 3.6 days, 95% CI = 1.9 to 5, range = 1-8 days) than the three donations that occurred in quartets during the same time period (latency = 6, 32, and 34 days). During controlled introduction trials, food sharing occurred in 6 of 11 possible cases between familiar bats in the quartets but only in 2 of 20 possible cases between unfamiliar bats in those same quartets (OR = 0.09, df = 1, p = 0.012).

*5. Relationships appeared to develop faster during controlled introductions*

During the controlled introductions trials, first donations were observed on average 33 days post-introduction (95% CI = 1 to 56, range = 1 to 193, n = 12 dyads) and first grooming was observed on average 24 days post-introduction (95% CI = 0 to 43, range = 1 to 205, n = 23 dyads). During the mixed-group trials, first donations in new dyads were observed on average 247 days after their introduction

(95% CI = 227 to 267, range = 66 to 556 days, n = 83 dyads, Figure S2). The first evidence of new grooming was seen on average 198 days after their introduction (95% CI = 186 to 209, range = 7 to 546, n = 351 dyads, Figure S2). During the mixed-group trials, the appearance of first donations became more probable over time (OR = 1.56, n = 3072, p = 0.0099), so new sharing relationships appeared to form gradually (Figures S2, S3).

*6. New grooming preceded new food sharing more than expected by chance*

It is important to note that our tests of whether new grooming occurred before new food sharing are highly conservative (i.e. biased away from detecting new grooming before new food sharing), because the actual first grooming events in a new pair almost certainly occurred before our first observations of it, whereas the first food donations we observed were likely to be the actual first donations. Food donations were only necessary during the 1-hour trial when we observed them. Bats were only focal sampled during fasting trials, and they were only in need during the fasting trials, because we isolated and fed them immediately after every trial. In contrast, grooming between the same bats could occur at any time during the days before the same dyad was sampled again (median gap period = 8 days, inter-quartile range = 5 to 14 days). In sum, we sampled close to 100% of the time when food sharing was necessary, but less than 2% of the time when grooming could have occurred. Additionally, although fasting trials increase the probability the subject will receive food, they also decrease the probability the subject will groom others (see '10. Grooming Symmetry' below). Therefore, when we observed the first grooming and sharing events during the same fasting trial, it is very likely that the first grooming actually occurred in the days before this trial.

Despite this conservative bias, we still observed new grooming events before new sharing more than expected based on their relative frequencies. In 57% of 87 new pairs, we observed grooming (but not sharing) during fasting trials on days before the trial with the first food sharing, showing that grooming preceded sharing. In 32% of these pairs, we observed the first grooming and sharing during the same trial, also suggesting that grooming preceded sharing. In the remaining 9 pairs, we observed the first new donation without observing at least 5 seconds of grooming in a fasting trial.

Mutual grooming, i.e. both bats grooming each other, in a trial is a better indication than one-way grooming of relationship development. We observed mutual grooming before the first trial with food sharing in 40% of the new sharing pairs, which is more than twice what is expected from our null model in which we randomly swapped the labels of whether events were 'grooming' or 'sharing' ($p < 0.0001$, expected frequency = 12%, 95% CI = 9% to 15%).

*7. Grooming trajectories over time predicted new sharing*

The age composition of new potential relationships affected the pre-donation grooming rate trajectories. For adult past strangers, the grooming probabilities increased for all recipients, including those that never donated (OR = 1.12, p = 0.004), and they increased significantly faster for grooming recipients that later donated (OR = 1.49, $p < 0.0002$; interaction: OR = 1.45, p = 0.017, Figure 1b). For new potential relationships with captive-born bats, however, the grooming probabilities actually decreased for grooming recipients that never donated (OR = 0.90, p = 0.01), and they tended to increase for recipients that did later donate (OR = 1.33, p = 0.04; interaction: OR = 1.72, $p < 0.0001$), Figure 1c).

*8. Grooming before versus after new sharing*

Grooming increased before but not after first donations in new relationships. We did not detect a difference in this effect between adults and captive-born bats (three-way interaction: p=0.55). The same pattern (Fig. 2) was found in new relationships between adults (interaction: OR = 1.60, p = 0.013; before: OR = 1.49, p = 0.012; after: OR = 1.01, p = 0.45) and in new relationships with captive-born bats (interaction: OR = 1.45, p = 0.009; before: OR = 1.33, p = 0.014; after: OR = 1.06, p = 0.34).

*9. Reciprocal development of food sharing*

Among adult past strangers, the proportion of previous trials in which bat A fed B predicted the occurrence of the first new reciprocal donation from bat B to A (OR = 6.00, n = 235, p = 0.016), and the number of previously unfamiliar pairs that donated food in both directions during the study period was greater than expected if new donations were random (p = 0.0001, observed bidirectional pairs = 13, expected = 4.6, expected 95% CI = 1 to 9). Note that the probability of reciprocation is low because new food sharing rates were overall low, all bats had access to multiple donors, and most sharing occurred among familiar bats with established relationships.

*10. Grooming symmetry*

Previous studies of raising-the-stakes have focused on grooming symmetry within short time periods [4-9], but our experimental design did not allow for precise measures of grooming symmetry within each dyad for three main reasons. First, grooming symmetry was reduced within fasting trials. A fasted subject was twice as likely to be groomed by a groupmate (13% probability) than to groom a groupmate (6% probability) because the fasted bat was typically 'greeted' by many groupmates at the start of the trial, which involves receiving simultaneous one-way grooming from several bats, and the subject usually spent less time grooming and more time begging (trying to lick the mouth of a potential donor). Second, we know that grooming rates were increasing over time, were symmetrical across dyads (mantel test: $r = 0.77$, $p < 0.0002$), and were not sufficiently sampled to accurately estimate the true grooming rate within each dyad. Two under-sampled estimates of the same value will converge (i.e. appear more symmetrical) over time merely because greater sample sizes lead to more precise estimates of the two grooming rates. Third, any observed increase in grooming symmetry over time could be driven by age effects, because mutual grooming (and hence grooming symmetry) is lower when one bat is not yet an adult [10].

*11. Evidence that new grooming and sharing are not both caused by proximity*

One null hypothesis is that bats initiate new grooming and sharing based entirely on proximity, and the relationship between new grooming and new sharing is therefore spurious. If so, new grooming rates should correlate with new sharing when strangers were able to freely associate (during the mixed-group period), but when strangers were forced into close proximity (during the controlled introduction trials), then this correlation should be much smaller or disappear entirely, because we have removed most of the variation in proximity (as proximity was roughly equal between all the bats in the small cage). In other words, if variation in proximity is actually driving the correlation between grooming and sharing, then removing this variation (with forced close contact) should reveal the lack of an association between grooming and sharing. In sharp contrast to this prediction, the estimated effect of new grooming given on new food received was greater during the controlled introduction periods compared to the same effect during the mixed-group trials where proximity was allowed to vary (forced close proximity: OR = 5.44, $p = 0.037$; variable proximity: OR = 1.63, $p = 0.033$; network logistic regression in the sna R package).

*12. Evidence for 'raising the stakes' in chimpanzees*

Previous evidence for 'raising the stakes' in nonhuman social relationships came from observations of grooming among familiar male chimpanzees after the death of an alpha male (5). The authors suggested that, during this period of social instability, these groupmates may have needed to re-establish their relationships, and that a diminishing threat of violence led to the increasing rates of grooming (5). Although the increase in grooming rates is consistent with each male 'raising the stakes' to assess the risk of aggression from their grooming partner, it might have also resulted from a general decline in vigilance against possible aggression from any other groupmate.

*13. Evidence for contingent reciprocity in a long-term relationship*

Evidence for reciprocity is controversial because researchers debate about whether putative cases of reciprocity involve helping that is both actually conditional on past experience and costly enough to be exploited by cheating (11). Depending on the relative payoffs for actors and receivers (12), even strictly conditional cooperation might, in theory, represent risk-free cooperative acts that lead to byproduct benefits, i.e. pseudo-reciprocity (13). To show that cooperative investments are contingent on the recipient's previous behavior, one must prevent reciprocation of a natural form of costly cooperation and, in doing so, show that this has induced a subsequent decrease in the actor's costly investments. Clear contingency has been demonstrated in partners lacking long-term bonds (11), as in the resource exchange between plants and fungi (14), or between rats trained to pull food for partners (15-17), but it has yet to be experimentally demonstrated in food-sharing vampire bats (2-3) or in any long-term social relationship. This lack of evidence is not surprising because stable long-term relationships are, by definition, difficult to perturb.

In our opinion, no studies have clearly shown that non-reciprocation leads to reduced investments in the context of a long-term cooperative relationship, but many studies have come close and have several of these components (1-3,11,15-24). For example, flycatcher pairs preferentially mobbed with neighboring pairs that helped them mob previously (22-23), vervet monkeys received more grooming after their ability to provide food was experimentally elevated (20), and dwarf mongoose received more grooming after their perceived contributions to cooperative sentinel behavior were experimentally elevated by playbacks (21). These studies suggest that the actors preferentially invested in more cooperative partners, but interestingly, the cooperative return benefit in all three examples is a public good (i.e. mobbing, opening a food cache, and sentinel behavior), not a reciprocal investment directed back to the actor (like sharing food with a specific individual). For example, when food is shared automatically (as when opening a cache in the case of the vervet monkeys), being socially connected to the most successful hunter is likely to bring benefits to any given individual; however, when food is actively and voluntarily shared with specific individuals (as when regurgitated), then that connection is only as beneficial as the hunter's propensity to share food with that specific individual over others. In such cases, when cooperative investments are highly variable and directed to specific partners, one must explain how animals might identify what cooperative partnerships are most beneficial for them specifically. The raising-the-stakes hypothesis does exactly this by proposing that returns on small cooperative investments are used to predict returns on larger investments.

### SI Figures and Tables

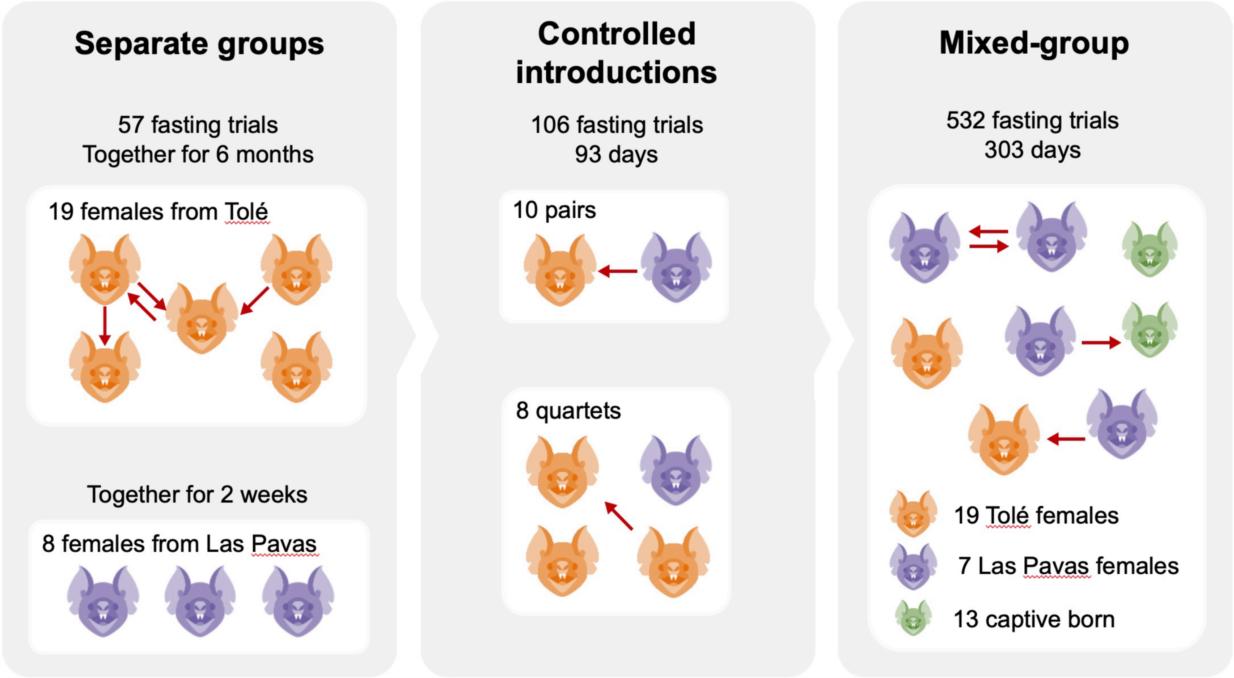


**Figure S1 | Experiment overview**

To see how vampire bats form new social bonds, we created groups of bats from two different sites (colors), then we induced and sampled food sharing and grooming events between bats that are either previously familiar or unfamiliar. Red arrows depict food sharing events during repeated fasting trials. For details of controlled introductions, see SI Table 2. Icons from icons8.com used under a Linkware license.

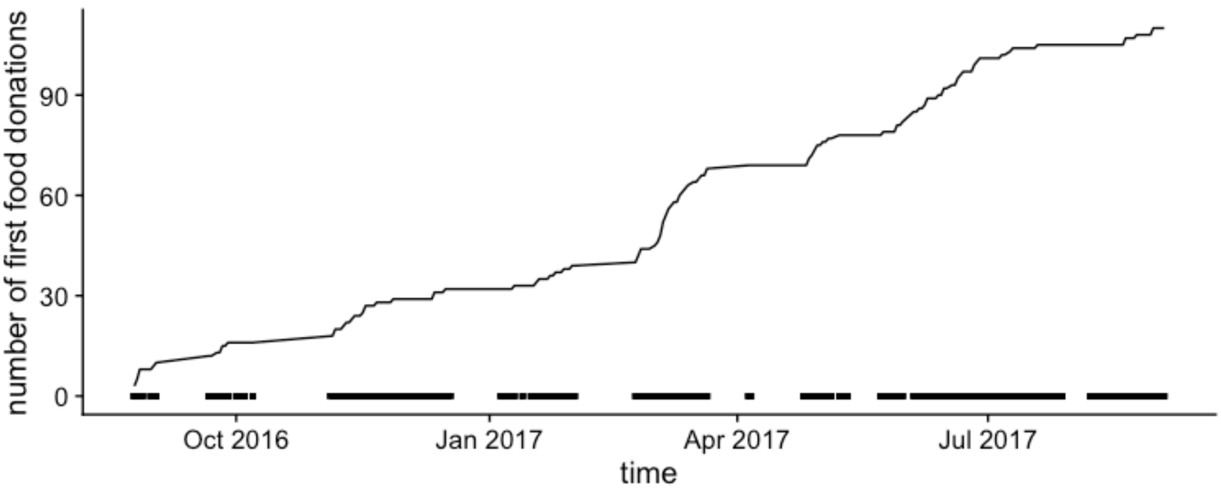


**Figure S2 | First food donations over time**
New food-sharing relationships accumulated gradually over time. Black rectangles above X-axis show the occurrence of fasting trials.

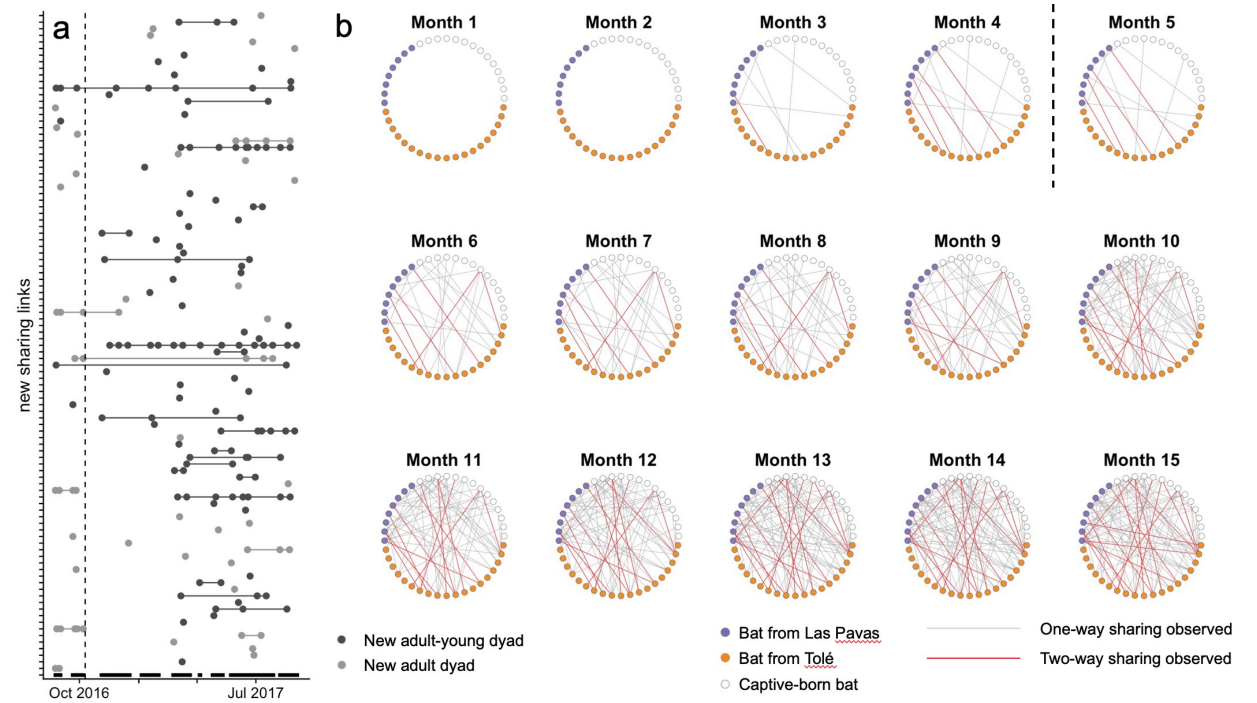


**Figure S3 | Gradual development of new food-sharing relationships**

Panel A shows food donations (points) over time (x-axis) within new actor-receiver relationships (y-axis) between two adult females (grey points) or with a captive-born bat (black points). Repeated dyadic donations are connected by horizontal lines. The end of the controlled introduction period, after which all bats could interact freely (months 1-4), is shown by the vertical dotted line. Black rectangles above the x-axis show the fasting trials, when new donations could be observed. Panel B shows the monthly formation of the food-sharing network between Las Pavas bats (orange), Tolé bats (purple), and captive-born bats (white). Grey edges show one-way sharing and red edges show two-way sharing. Two-way sharing occurred more often than expected by chance (see results).

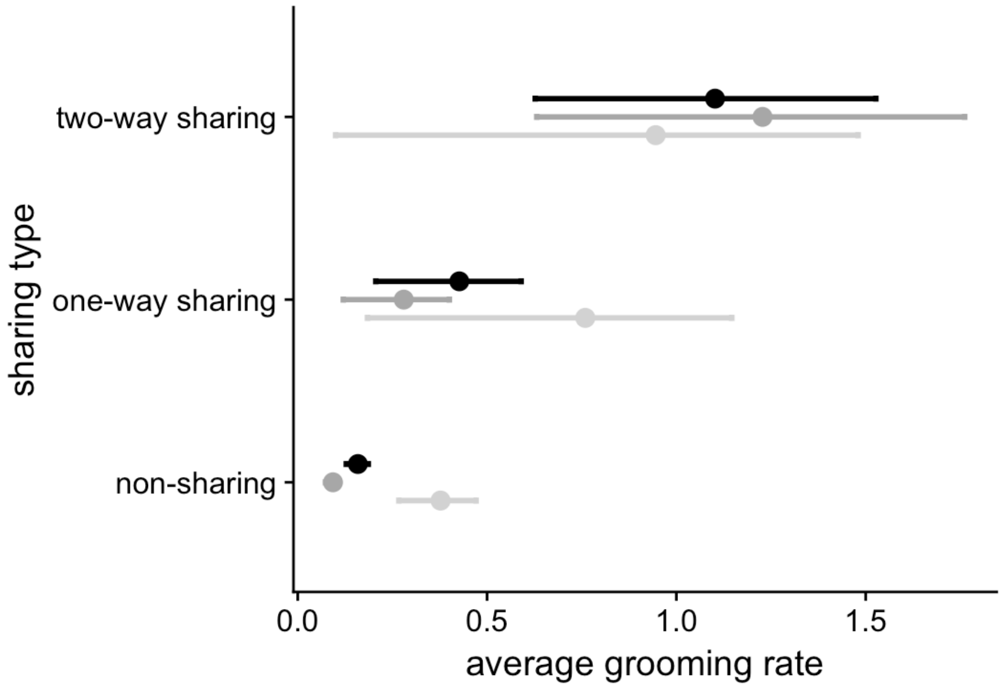


**Figure S4 | Dyadic grooming rates predict new food-sharing relationships.**
Mean within-dyad grooming rates, with bootstrapped 95% confidence intervals, are shown for three possible outcomes (y-axis) and for all potential relationships (black), potential relationships with captive-born bats (grey), and potential relationships between adult strangers (light grey).

**Table S1 | Glossary.**
Definition of terms used in the text.

| Term | Definition |
|---|---|
| Dyad | An undirected pair of bats (e.g. AB, BC, AC) |
| Potential relationship | A directed pair of actor and receiver bats (e.g. AB, BA, AC) |
| Relationship | A directed actor-receiver pair that is observed to groom or share food during fasting trials. |
| New relationship | Relationship between bats that first met during the experiment, excluding mother-offspring dyads. |
| Social bond | The unobserved underlying social relationship (as experienced by the animal) that we infer from observations. |

**Table S2 | Controlled introductions**
The same bats were used in multiple introductions. Bats were moved to and from groups to make new combinations or because of health issues (pregnancy, weight loss). Bats not in a small cage group during controlled introduction trials were kept with familiar individuals in a flight cage.

| No. | Group type | No. trials (range of days together) | Adult female bats (*Las Pavas stranger) | Opportunities for new sharing | Introduction date |
|---|---|---|---|---|---|
| 1 | quartet | 1 (1 day) | scs, hilga, rc, eve* | 3 | 2016.07.06 |
| 2 | quartet | 1 (1 day) | ccs, sss, sc, una* | 3 | 2016.07.06 |
| 3 | quartet | 1 (1 day) | scc, sd, c, dos* | 3 | 2016.07.06 |
| 4 | quartet | 1 (1 day) | csc, ss (w/pup), s, tes* | 3 | 2016.07.06 |
| 5 | pair | 1 (1 day) | ccc, cat* | 1 | 2016.07.06 |
| 6 | pair | 1 (1 day) | dcd, ivy* | 1 | 2016.07.06 |
| 7 | pair | 1 (1 day) | dd, six* | 1 | 2016.07.06 |
| 8 | pair | 1 (4 days) | d (w/pup), ola* (w/pup) | 1 | 2016.07.02 |
| 9 | quartet | 17 (1–44 days) | sd, scs, d (w/pup), una* | 32 | 2016.08.24 |
| 10a | quartet | 5 (1–9 days) | s, rc, hilga, dos* | 9 | 2016.08.24 |
| 10b | quartet | 12 (1–44 days) | s, rc, ccc (w/pup), dos* | 21 | 2016.09.21 |
| 11 | quartet | 17 (1–44 days) | ccs, sc, sss, tes* | 27 | 2016.08.24 |
| 12 | pair | 10 (1–44 days) | dd, cat* | 10 | 2016.08.24 |
| 13 | pair | 10 (1–44 days) | c, ivy* | 10 | 2016.08.24 |
| 14 | pair | 5 (1–10 days) | csc, six* | 5 | 2016.08.24 |
| 15 | pair | 9 (1–44 days) | dcd, eve | 9 | 2016.08.24 |
| 16 | pair | 9 (1–97 days) | ss (w/pup), ola* (w/pup) | 19 | 2016.08.24 |
| 17 | pair | 4 (1–7 days) | cd, six* | 4 | 2016.09.21 |

**Data**
**Dataset S1.** genotypes.csv
Microsatellite genotypes used to assess relatedness.
**Dataset S2.** vampire_maternal_kinship.csv
Maternal pedigree data
**Dataset S3.** new_bonds_data.Rdata
Food sharing and allogrooming data
**Dataset S4.** new_bonds_analysis22.R
R script for analyzing data
